# Epigenetic silencing of miRNA-338-5p and miRNA-421 drives SPINK1-positive prostate cancer

**DOI:** 10.1101/376277

**Authors:** Vipul Bhatia, Anjali Yadav, Ritika Tiwari, Shivansh Nigam, Sakshi Goel, Shannon Carskadon, Nilesh Gupta, Apul Goel, Nallasivam Palanisamy, Bushra Ateeq

## Abstract

**Purpose:** Serine Peptidase Inhibitor, Kazal type-1 (SPINK1) overexpression defines the second most recurrent and aggressive prostate cancer (PCa) subtype. However, the underlying molecular mechanism and pathobiology of SPINK1 in PCa remains largely unknown.

**Experimental Design:** MicroRNA-prediction tools were employed to examine the *SPINK1-*3’UTR for miRNAs binding. Luciferase reporter assays were performed to confirm the *SPINK1-*3’UTR binding of shortlisted miR-338-5p/miR-421. Further, miR-338-5p/-421 overexpressing cancer cells (SPINK1-positive) were evaluated for oncogenic properties using cell-based functional assays and mice xenograft model. Global gene expression profiling was performed to unravel the biological pathways altered by miR-338-5p/-421. Immunohistochemistry and RNA *in-situ* hybridization was carried-out on PCa patients’ tissue microarray for SPINK1 and *EZH2* expression respectively. Chromatin immunoprecipitation assay was performed to examine EZH2 occupancy on the miR-338-5p/-421 regulatory regions. Bisulfite sequencing and methylated DNA-immunoprecipitation was performed on PCa cell lines and patients’ specimens.

**Results:** We established a critical role of miRNA-338-5p/-421 in post-transcriptional regulation of *SPINK1*. Ectopic expression of miRNA-338-5p/-421 in SPINK1-positive PCa cells abrogate oncogenic properties including cell-cycle progression, stemness and drug resistance, and show reduced tumor burden and distant metastases in mice model. Importantly, we show SPINK1-positive PCa patients exhibit increased EZH2 expression, suggesting its role in miRNA-338-5p/-421 epigenetic silencing. Furthermore, presence of CpG dinucleotide DNA methylation marks on the regulatory regions of miR-338-5p/-421 in SPINK1-positive PCa cells and patients’ specimens confirms epigenetic silencing.

**Conclusion:** Our findings revealed that miRNA-338-5p/-421 are epigenetically silenced in SPINK1-positive PCa, while restoring the expression of these miRNAs using epigenetic drugs or synthetic mimics could abrogate SPINK1-mediated oncogenesis.

**TRANSLATIONAL IMPACT:** We establish a regulatory model involving the functional interplay between SPINK1, miRNA-338-5p/miRNA-421 and EZH2, thereby, revealing hitherto unknown mechanism of SPINK1 up-regulation in SPINK1-positive subtype. Our findings provide a strong rationale for the development of potential therapeutic strategies for SPINK1-positive malignancies. We demonstrate that restoring miRNA-338-5p/miRNA-421 expression using epigenetic drugs including DNMTs inhibitors in combination with HDACs or HKMTs inhibitors or miRNA synthetic mimics in SPINK1-positive prostate cancer abrogate SPINK1-mediated oncogenicity. The major findings of this study will not only advance the prostate cancer field, but will also be valuable for treatment and disease management of other SPINK1-positive malignancies.

## Introduction

Prostate Cancer (PCa) is characterized by extensive molecular heterogeneity and varied clinical outcomes (1). Multiple molecular subtypes involving recurrent genetic rearrangements, DNA copy number alterations, and somatic mutations have been associated with this disease (1-3). Majority of the PCa patients harbor gene rearrangements between members of the E26 transformation specific (*ETS*) transcription factor family and the androgen-regulated transmembrane protease serine 2 (*TMPRSS2*), most recurrent (~50%) being *TMPRSS2-ERG*, a gene fusion involving the v-ets erythroblastosis virus E26 oncogene homolog (*ERG*) (3,4). The ERG transcription factor encoded by *TMPRSS2-ERG* fusion is known to drive cell invasion and metastases, induce DNA damage *in-vitro* and focal pre-cancerous prostatic intraepithelial neoplasia (PIN) lesions in transgenic mice (5).

While *TMPRSS2-ERG* fusion forms the most frequent molecular subtype, a significant subset of *ETS*-negative (–) PCa show overexpression of Serine Peptidase Inhibitor, Kazal type-1 (*SPINK1*) in ~10-15% of the total PCa patients, a distinct subtype defined by overall higher Gleason score, shorter progression-free survival and biochemical recurrence (6,7). SPINK1 promotes cell proliferation and invasion through autocrine/paracrine signaling and mediate its oncogenic effects in part through EGFR interaction by activating downstream signaling. Monoclonal antibody against EGFR showed only a marginal decrease in the growth of SPINK1-positive (+) xenografts in mice, supporting involvement of EGFR-independent oncogenic pathways (8).

Although, genomic events such as genetic rearrangements and somatic mutations constitute most recurrent oncogenic aberrations, many could also be attributed to epigenetic alterations. Earlier studies have shown that aberrant expression of Enhancer of Zeste Homolog 2 (EZH2) owing to genomic loss of miRNA-101 (9) or hypermethylation of miR-26a (10) constitutes a common mechanism across several solid cancers including prostate. EZH2, being a key component of the Polycomb-Repressive Complex 2 (PRC2) mediates trimethylation on the histone 3 lysine 27 (H3K27me3), leading to gene silencing (11). However, phosphorylated form of EZH2 is known to switch its function from Polycomb repressor to transcriptional coactivator of androgen receptor in castration-resistant prostate cancers (CRPC) (12). Moreover, recent studies have shown PRC2 epigenetically suppresses the expression of several tumor suppressive miRNAs such as, miR-181a/b, miR-200b/c, and miR-203, while these miRNAs in turn directly target PRC1 members, namely *BMI1* and *RING2*, in breast and prostate cancer thereby reinforcing the repressive molecular circuitry (13).

Although SPINK1+ subtype forms a well-defined and second most prevalent subset of PCa (4,14), the underlying mechanism involved in its upregulation is poorly understood and remains a matter of conjecture. Further, overexpression of *SPINK1* is not ascribed to chromosomal rearrangement, deletion, or amplification (14), and thus alludes to a possible transcriptional or post-transcriptional regulation. A recent study showed the transcriptional activation of *SPINK1* along with gastrointestinal (GI) lineage signature genes in CRPC patients (15). Our study focuses on the post-transcriptional regulation of *SPINK1* expression by miR-338-5p and miR-421, which are epigenetically silenced in SPINK1-positive prostate cancer. We also provide evidence that EZH2 acts as an epigenetic switch, thereby promoting transcriptional silencing of miR-338-5p/miR-421 by establishing H3K27me3 repressive marks, thus leading to SPINK1 overexpression. Collectively, our findings suggest potential benefits with epigenetic inhibitors or synthetic miR-338-5p/-421 mimics as adjuvant therapy for the treatment of aggressive *SPINK1+* malignancies.

## Materials and Methods

### Human Prostate Cancer Specimens

All prostate cancer (PCa) specimens used in this study were procured from King George’s Medical University, Lucknow, India. Clinical specimens were collected after obtaining written informed consent from the patients and Institutional Review Board approvals from the King George’s Medical University, Lucknow and Indian Institute of Technology, Kanpur, India. A total of 20 PCa specimens were selected for this study based on the *SPINK1* and *TMPRSS2-ERG* status, confirmed by qPCR, immunohistochemistry and Fluorescent in situ hybridization (FISH) for confirming gene rearrangement (4). Prostate cancer tissue microarrays (TMA) specimens (n=238) were obtained from Dept. of Pathology, Henry Ford Health System, Detroit, Michigan, USA, after getting written informed consent from the patients and approval from Institutional Review Board. TMAs were stained for SPINK1 and *EZH2* by performing immunohistochemistry (IHC) and RNA *in situ* hybridization (RNAISH) respectively.

### Gene expression array analysis

For global gene expression profiling, total RNA was isolated from stable 22RV1-miR-338-5p, 22RV1-miR-421 and 22RV1-CTL cells and subjected to Agilent Whole Human Genome Oligo Microarray profiling (dual color) using Agilent platform (8×60K format) according to the manufacturer’s protocol. A total of three microarray hybridizations were performed using each stable miRNA overexpressing cell line samples against control cells. Microarray data was normalized by following locally weighted linear regression (also known as Lowess). Differentially regulated genes were clustered using hierarchical clustering based on Pearson coefficient correlation algorithm to identify significant gene expression patterns. Further, multiple hypotheses testing adjustments were applied using Benjamini and Hochberg procedure to calculate the FDR-corrected *P*-values (with FDR< 0.05) for the differentially expressed genes. Data was filtered to include only features with significant differential expression (log_2_ fold change greater than 0.6 or less than -0.6, *P*< 0.05) i.e. ~1.6-fold average over- or under-expressed genes, were then used for the deregulated biological processes using DAVID bioinformatics platform. Further, molecular signatures that were enriched upon miRNA overexpression with respect to control were analyzed using Gene set enrichment analysis (GSEA). A network-based analysis of critical miR-338-5p/-421 overlapping biological pathways was generated using Enrichment Map, a plug-in for Cytoscape network visualization software (http://baderlab.org/Software/EnrichmentMap/).

### Data Mining and Computational Analyses

#### Integrative MicroRNA target Prediction

MicroRNA prediction programs, namely miRanda, miRMap, PITA and RNA Hybrid were used to predict miRNAs targeting 3’UTR of *SPINK1* (Fig. 1A and Supplementary Table S1). The correlation between the expression of the predicted miRNAs and *SPINK1* was analyzed by employing RNA-Seq data for the TCGA-PRAD cohort. For Fig. 1A (lower panel) and Supplementary Table S6 miRanda was used to predict the putative binding sites of the miR-338-5p/-421 on the 3’UTR of target genes.

**Figure 1.**
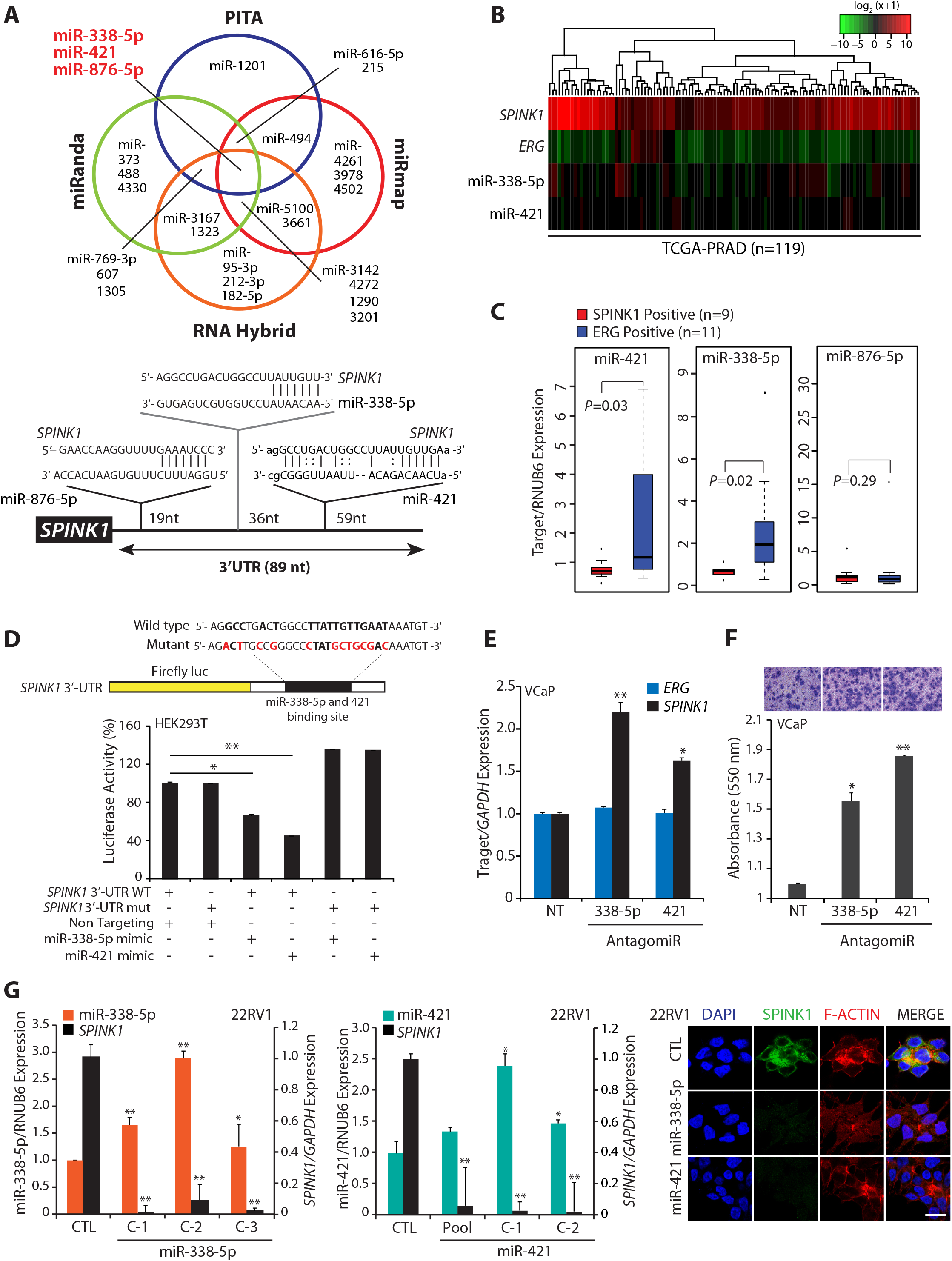
MiR-338-5p and miR-421 are differentially expressed in *SPINK1+/ERG-*fusion-negative prostate cancer. **(A)** Venn diagram displaying miRNAs computationally predicted to target *SPINK1* by PITA, miRmap, miRanda and RNAHybrid (top panel). Schematic of predicted miR-338-5p, miR-421 and miR-876-5p binding sites on the 3′-UTR of *SPINK1* (bottom panel). **(B)** Heatmap depicting miR-338-5p and miR-421 expression in the *SPINK1*+/*ERG-*negative patients’ (n=119) in TCGA-PRAD dataset. Shades of red and green represents log_2_ (x+1), where x represents the gene expression value. **(C)** Taqman assay showing relative expression for miR-338-5p, miR-421 and miR-876-5p in *SPINK1+* and *ERG+* PCa patients’ specimens (n=20). Data represents normalized expression values with respect to RNUB6 control. Error bars represent mean ± SEM. *P-*values were calculated using two-tailed unpaired Student’s *t* test. **(D)** Schematic of luciferase reporter construct with the wild-type or mutated (altered residues in red) *SPINK1* 3’ untranslated region (3’UTR) downstream of the Firefly luciferase reporter gene (top panel). Luciferase reporter activity in HEK293T cells co-transfected with wild-type or mutant 3′-UTR *SPINK1* constructs with mimics for miR-338-5p or miR-421. **(E)** QPCR data showing *SPINK1* and *ERG* expression in VCaP cells transfected with antagomiRs as indicated (n=3 biologically independent samples; data represent mean ± SEM). **(F)** Boyden chamber Matrigel invasion assay using same cells as in (E). Representative fields of the invaded cells are shown in the inset. **(G)** QPCR analysis demonstrating *SPINK1* and miRNAs expression in stable 22RV1-miR-338-5p (left panel) and 22RV1-miR-421 cells (middle panel) (n=3 biologically independent samples; data represent mean ± SEM). Immunostaining for SPINK1 (right panel). Scale bar represents 20µm. For all panels ^∗^*P* ≤ 0.05 and ^∗∗^*P* ≤ 0.005 using two-tailed unpaired Student’s *t* test.

#### Integrative analyses for The Cancer Genome Atlas Prostate Adenocarcinoma (TCGAPRAD) data

For gene association studies between miRs-338-5p, -421, miR-876-5p, *SPINK1*, *ERG*, and *EZH2* Illumina HiSeq mRNA and miRNA-Seq data along with clinical information was downloaded from TCGA-PRAD dataset. Overexpression of SPINK1 in PCa exhibits outlier-expression in ~10-15% of the total PCa cases (6). Thus, to stratify patients with increased expression of *SPINK1*, we sorted TCGA patients’ samples on the basis of increasing *SPINK1* expression (descending order), and divided the dataset into four equal parts by employing Quartile-based normalization method (16), the top 25% of the patients (n=119) corresponding to the upper quartile (QU, log_2_ (RPM+1)>5.468 or log_2_ (normalized count+1)>1.892), were assigned as *SPINK1* high or *SPINK1*+ patient samples and the lower quartile (QL, log_2_ (RPM+1)<1.124 or log_2_ (normalized count+1)<-2.611), were considered as *SPINK1* low or *SPINK1*-negative samples. Also, we found about 18 patients with outlier expression of *SPINK1* with log_2_ (RPM+1) of greater than 11.984, which were included in the heat map representation of *SPINK1* positive TCGA patients in Fig. 1B. No further cut-offs were applied for miR-338-5p, miR-421 and *ERG* expression, corresponding expression values (based on *SPINK1* cut-off) were considered for these genes for further analysis. Hierarchical Clustering of miRs-338-5p, -421 or 876-5p, *SPINK1*, and *ERG* were employed using heatmap.2 of R’s gplot package.

For Kaplan-Meier survival analysis, the survival data included sample type (primary tumors), days to first biochemical recurrence and days to last follow-up for TCGA-PRAD patients was considered. The samples were divided into two groups, higher and lower miRNA expression groups according to the expression level of a miR-338-5p/-421 using Cox proportional hazards regression model in R. Next, we performed a 13-year survival analysis of these miRNAs using Kaplan-Meier survival analysis (17) by employing survival package (https://cran.r-project.org/web/packages/survival), and statistical significance was computed using the log-rank test. For clinical relevance of miR-338-5p/-421, TCGA-PRAD dataset was analyzed for the association of these miRNAs with clinical parameters such as primary Gleason score. Data analysis was performed by one-way analysis of variance with Tukey’s post hoc test for multiple comparisons, and student’s *t*-test was applied for comparison between two groups.

The MSKCC cohort (Cancer Cell, 2010) data was retrieved from cBioPortal (http://www.cbioportal.org/) for *SPINK1* and *EZH2* expression in the prostate cancer patients (n=85), and oncoprints were generated using default parameters (mRNA expression z-score threshold ±2 vs normal). Further, to ascertain possible association between *EZH2*, *SPINK1* and miR-338-5p/-421, TCGA patients’ samples were stratified on the basis of increasing *EZH2* expression, and divided the dataset into four equal quartiles, the top 25% of the patients (n=119) corresponding to the upper quartile (QU, log_2_ (normalized count+1) >7.313), were considered as *EZH2* high patient samples and the lower quartile log_2_ (normalized count+1) <6.36), were considered as *EZH2* low samples. The corresponding expression values for *SPINK1*, miR-338-5p and miR-421 in *EZH2* high and *EZH2* low groups (without further cut-offs) were considered to association with *EZH2* expression (related to Fig. 5B).

### Statistical analysis

Statistical significance was determined by either two-tailed Student’s *t* test for independent samples or one-way Analysis of Variance (ANOVA), otherwise specified. The differences between the experimental groups were considered significant if the *p*-value of less than 0.05 was obtained. Error bars represent mean ± SEM. All experiments were repeated three times in triplicates.

### Supplementary methods

All further information can be found in the Supplementary Methods section.

## Results

### Identification of differentially expressed miRNAs in *SPINK1+/ERG*-fusion-negative prostate cancer

We employed four miRNA prediction algorithms, namely PITA (omicstools.com), miRmap (mirmap.ezlab.org), miRanda (microRNA.org) and RNAHybrid (BiBiserv2-RNAhybrid) to examine putative binding of miRNAs to the 3’ untranslated region (3’UTR) of *SPINK1* transcript. Notably, three miRs -338-5p, -421 and -876-5p were predicted as strong candidates by all four algorithms (Fig. 1A and Supplementary Table S1), and were taken forward for further investigation. To examine the differential expression of these three miRNAs between *SPINK1+* and *ERG*+ PCa patients’ specimens, RNA-seq data available at public repository, TCGA-PRAD was analyzed. Interestingly, hierarchical clustering of TCGA-PRAD RNA-Seq dataset exhibit reduced expression of miR-338-5p and miR-421 (miR-338-5p/-421) in *SPINK1+/ERG*-negative patient specimens (Fig. 1B). To validate further, we examined the expression of miR-338-5p/-421 and miR-876-5p in our PCa patients’ specimens. A significant lower expression of miR-338-5p and miR-421 was observed specifically in *SPINK1+* as compared to *ERG+* specimens, while no difference in miR-876-5p expression was noticed (Fig. 1C). To understand the clinical significance of miR-338-5p/-421, we stratified TCGA-PRAD patients’ data into high and low miRNAs expressing groups. Intriguingly, patients with low miR-338-5p expression showed significant (*p*=0.0024) association with decreased survival probability compared to high miRNAs group, while no such association was found in case of miR-421 (Supplementary Fig. S1A). Moreover, lower expression of miR-338-5p also associate with higher Gleason score (Supplementary Fig. S1B). An association of higher Gleason score with lower expression of both miRNAs was further confirmed in another independent cohort (GSE45604) (Supplementary Fig. S1C). In summary, *SPINK1+* subtype show lower expression of miR-338-5p/-421, which strongly associate with overall poor survival and aggressiveness of the disease.

### MiR-338-5p and miR-421 directly target *SPINK1* and modulate its expression

Having established an association between miR-338-5p/ -421 and *SPINK1* expression in PCa specimens (Fig. 1B, C), we next examined the ability of these miRNAs to bind to the 3’-untranslated region (3’UTR) of *SPINK1*. The wild-type (3’-UTR-WT) and mutant (3’-UTR-mut) *SPINK1* 3’-UTR cloned in Firefly/Renilla dual-luciferase reporter vectors were cotransfected with synthetic mimics for miR-338-5p or miR-421 in HEK293T cells, a significant reduction in the luciferase activity was noted with 3’-UTR-WT, while 3’-UTR-mut constructs failed to show any suppressive effect (Fig. 1D). We next evaluated the expression of these miRNAs in various PCa cell lines including 22RV1 (*SPINK1+*), *ETS-*fusion positive VCaP (*TMPRSS2-ERG*+) and LNCaP (*ETV1*+) cells. Supporting our observation in clinical specimens the cell line data also showed lower expression of miR-338-5p/-421 in the 22RV1 cells relative to fusion-positive cell lines (Supplementary Fig. S1D). To further ascertain that miR-338-5p/miR-421 specifically regulates *SPINK1*, we used antagomiRs to abrogate miR-338-5p and miR-421 expression (anti-338-5p and anti-421, respectively) in VCaP cells (Supplementary Fig. S1E). As expected, anti-338-5p or anti-421 significantly induced *SPINK1* expression in VCaP cells with concomitant increase in cell invasion and migration (Fig. 1E, F and Supplementary Fig. S1F, G), while there was no change in the endogenous ERG expression (Fig. 1E and Supplementary Fig. S1F). Conversely, we observed that 22RV1 cells stably overexpressing miR-338-5p or miR-421 (22RV1-miR-338-5p and 22RV1-miR-421, respectively) show a significant reduction in *SPINK1* expression at both transcript (~80-90%) and protein (Fig. 1G) levels. Since, SPINK1 overexpression has also been implicated in colorectal, lung, pancreatic, breast and ovarian cancers (18,19), we sought to examine if SPINK1 is regulated by a similar mechanism in cancers of different cellular/tissue origins. Thus, we determined the status of SPINK1 expression in multiple cancer cell lines (Supplementary Fig. S2A, B). Further, *SPINK1+* cancer cell lines, namely, colorectal (WiDr), melanoma (SK-MEL-173), pancreatic (CAPAN-1) and prostate (22RV1) upon transfecting with mimics for miR-338-5p or miR-421 showed a significant decrease in SPINK1 expression both at transcript and protein levels (Supplementary Fig. S2C, D). This provides irrevocable evidence that these two miRNAs modulate the expression of *SPINK1* transcript irrespective of the tissue background. Furthermore, to ascertain whether decrease in oncogenic properties is indeed due to miR-338/-421 mediated reduction in SPINK1 expression, a rescue cell migration assay using human recombinant SPINK1 (rSPINK1) was performed. As expected, miR-338 and miR-421 overexpressing 22RV1 cells show decrease in cell migration, while adding rSPINK1 to these miRNAs overexpressing cells rescued the invasive phenotype, indicating that miR-338/-421 mediated effects are indeed due to decrease in SPINK1 expression (Supplementary Fig. S2E).

### Ectopic expression of miR-338-5p and miR-421 attenuate SPINK1-mediated oncogenesis

SPINK1 overexpression is known to contribute to cell proliferation, invasion, motility and distant metastases (4,14,20). Hence, to understand the functional relevance of miR-338-5p/ -421, we examined 22RV1-miR-338-5p and 22RV1-miR-421 stable cells for any change in their oncogenic properties. Both 22RV1-miR-338-5p (C1 and C2) and 22RV1-miR-421 (pooled and C1) cells showed a significant decrease in cell proliferation compared to control (22RV1-CTL) cells (Fig. 2A). Similarly, a significant reduction in invasive properties of 22RV1-miR-338-5p and 22RV1-miR-421 cells were noted (~40% and 60% respectively) (Fig. 2B). While, only a modest decrease in cell proliferation and invasion was observed in pooled 22RV1-miR-338-5p cells (Supplementary Fig. S3A-C). To assess neoplastic transformation, soft agar colony formation assay was performed, where both 22RV1-miR-338-5p and 22RV1-miR-421 cells exhibited marked reduction (~60% and ~80% respectively) in number and size of the colonies (Fig. 2C). Likewise, 22RV1-miR-338-5p and 22RV1-miR-421 cells demonstrate significantly lower numbers (~70% and ~60% respectively) of dense foci (Fig. 2D). While, overexpressing miR-338-5p/-421 in immortalized benign prostate epithelial RWPE-1 cells show no significant change in cell proliferation or migration with miR-338-5p mimics transfection; conversely, RWPE-1 cells transfected with miR-421 showed marginal decrease in proliferation and migration (Supplementary Fig. S3D) Further, to demonstrate that miR-338-5p/-421 modulate *SPINK1* expression and attenuate SPINK1-mediated oncogenicity irrespective of the tissue background, we performed functional assays using colorectal carcinoma WiDr cells (SPINK1+) stably overexpressing these miRNAs. As anticipated, a significant decrease in the oncogenic potential of the miR-338-5p/-421 overexpressing WiDr cells was observed (Supplementary Fig. S3E, F).

**Figure 2.**
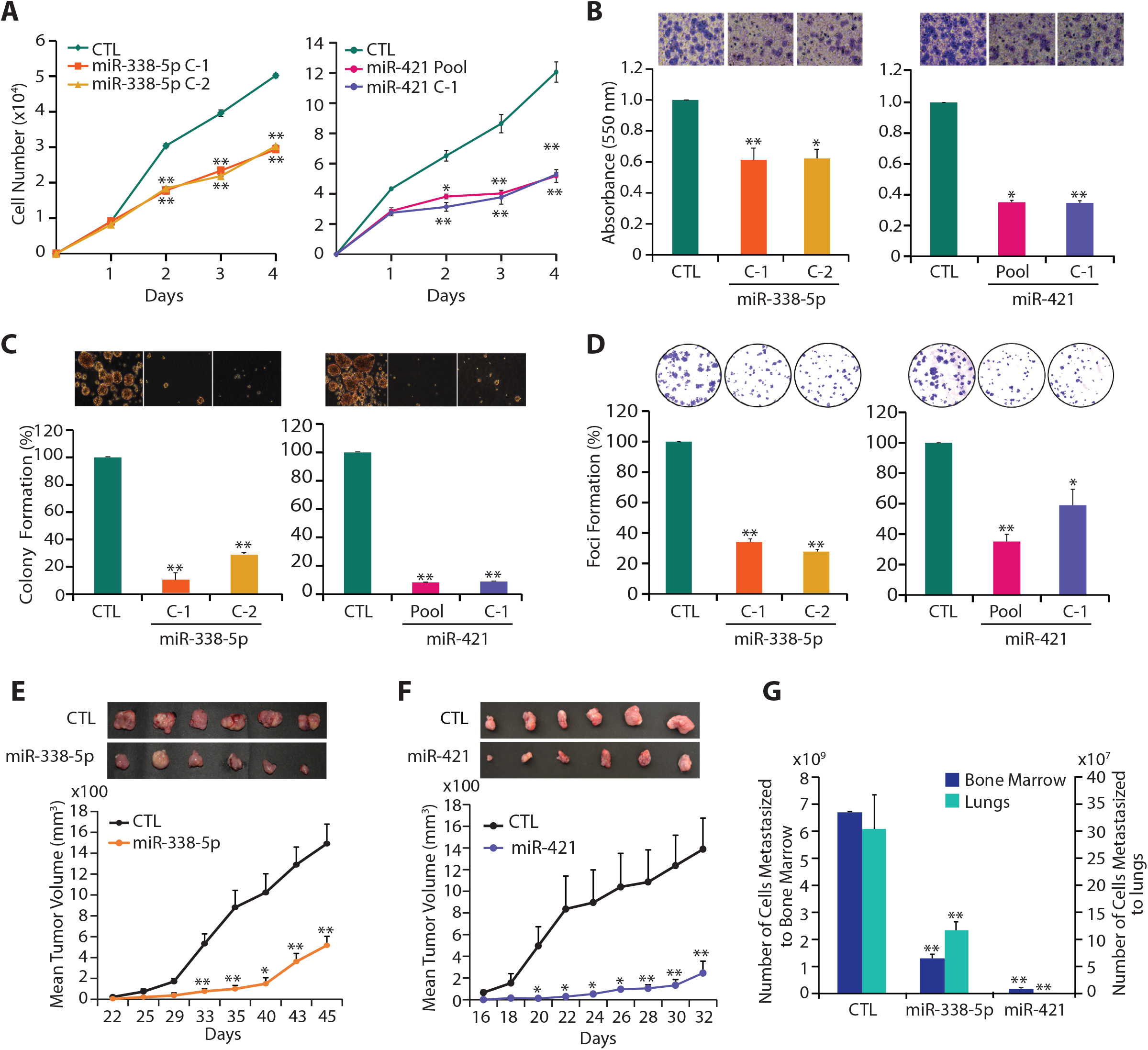
MiR-338-5p and miR-421 abrogates oncogenic properties of SPINK1-positive prostate cancer cells. **(A)** Cell proliferation assay using 22RV1-miR-338-5p, 22RV1-miR-421 and 22RV1-CTL cells at the indicated time points. **(B)** Boyden chamber Matrigel invasion assay using same cells as in (A). Representative fields with invaded cells are shown in the inset (n=3 biologically independent samples; data represent mean ± SEM). **(C)** Soft agar assay for anchorage-independent growth using same cells as in (A). Representative soft agar colonies are shown in the inset (n=3 biologically independent samples; data represent mean ± SEM). **(D)** Foci formation assay using same cells as in (A). Representative images depicting foci are shown in the inset (n=3 biologically independent samples; data represent mean ± SEM). **(E)** Mean tumor growth in NOD/SCID mice (n=8) subcutaneously implanted with stable 22RV1-miR-338-5p and 22RV1-CTL cells. **(F)** Same as (E), except stable 22RV1-miR-421 cells were implanted. **(G)** Same as (E and F), except genomic DNA extracted from the lung and bone marrow of the xenografted mice. Data represent mean ± SEM. ∗*P*≤ 0.05 and ^∗∗^*P*≤ 0.005 using two-tailed unpaired Student’s *t* test.

To examine tumorigenic potential of 22RV1-miR-338-5p and 22RV1-miR-421 cells *in-vivo*, chick chorioallantoic membrane (CAM) assay was performed, and relative number of intravasated cancer cells was analyzed. Consistent with *in-vitro* results, 22RV1-miR-338-5p and 22RV1-miR-421 cells showed significant reduction in the number of intravasated cells compared to control (Supplementary Fig. S4A, B). Likewise, a significant reduction in the tumor weight was recorded in the groups implanted with 22RV1-miR-338-5p and 22RV1-miR-421 cells (Supplementary Fig. S4C). To evaluate distant metastases, lungs and liver excised from the chick-embryos were characterized for the metastasized cancer cells. The groups implanted with miRNAs overexpressing cells revealed ~80% reduction in cancer cell metastases to lungs (Supplementary Fig. S4D), while no sign of liver metastases was observed in either group. Further, tumor xenograft experiment was recapitulated in immunodeficient NOD/SCID mice (n=8 per group) by subcutaneously implanting 22RV1-miR-338-5p, 22RV1-miR-421 and control 22RV1 cells into flank region, and trend of tumor growth was recorded. A significant reduction in the tumor burden was observed in the mice bearing miR-338-5p and miR-421 overexpressing xenografts as compared to control (~70% and 85% reduction respectively) (Fig. 2E, F). To examine spontaneous metastases, lung, liver and bone marrow specimens were excised from the xenografted mice, and genomic DNA was quantified for the presence of human specific *Alu*-sequences. A significant decrease (~85% for miR-338-5p and ~90% for miR-421) in cancer cell metastases was observed in the group implanted with miRNAs overexpressing cells (Fig. 2G). Similar to CAM assay, cancer cells failed to metastasize to murine liver (data not shown). Furthermore, a significant drop (~50%) in Ki-67-positive cells in the miRNAs overexpressing xenografts confirms that tumor regression was indeed due to decline in cell proliferation (Supplementary Fig. S4E). Taken together, our findings indicate that miR-338-5p/-421 downregulate the expression of *SPINK1* and abrogate SPINK1-mediated oncogenic properties and tumorigenesis.

### MiR-338-5p and miR-421 exhibit functional pleiotropy by regulating diverse biological processes

To explore critical biological pathways involved in the tumor-suppressive properties rendered by miR-338-5p/-421 in SPINK1+ cancers, we determined global gene expression profiles of miRNAs overexpressing 22RV1 cells. Our analysis revealed 2,801 and 2,979 genes significantly dysregulated in 22RV1-miR-338-5p and 22RV1-miR-421 cells respectively relative to control (log_2_ fold change of 0.6, *FDR*<0.05 and *p*<0.05) (Supplementary Table S2 and S3). Remarkably, ~22% (704 genes) of the downregulated and ~15% (506 genes) of the upregulated transcripts show an overlap in miR-338-5p and miR-421 overexpressing cells (90% confidence interval) (Fig. 3A), indicating that these two miRNAs regulate a significant number of common gene-sets and cellular processes. To examine biological processes commonly regulated by miR-338-5p/-421, we employed DAVID (Database for Annotation, Visualization and Integrated Discovery) and GSEA (Gene set enrichment analysis). Most of the downregulated genes were associated with DNA double-strand break repair by homologous recombination, cell cycle regulation including G2/M-phase transition, stem-cell maintenance, histone methylation and negative regulation of cell-cell adhesion. Whereas, genes involved in negative regulation of gene expression or epigenetics, intrinsic apoptotic signaling pathways, negative regulation of metabolic process and cell cycle were significantly upregulated (Fig. 3B, Supplementary Table S4 and S5). Moreover, GSEA also revealed enrichment of gene signatures associated with oncogenic pathways and cancer hallmarks. Conversely, 22RV1-CTL cells showed significant enrichment of genes involved in sustaining proliferative signaling (EGFR and MEK/ERK) and cell cycle regulators (E2F targets and G2/M transition). While, positive enrichment for tumor suppressive p53 signaling was found in miRNA overexpressing cells as compared to control (Fig. 3C), indicating its role in reduced oncogenicity. Additionally, an overlapping network of pathways using Enrichment map revealed regulation of cell-cycle phase transition and DNA repair pathways (overlap coefficient=0.8, *p*<0.001, *FDR*=0.01), as one of the significantly enriched pathways for both miRNAs (Supplementary Fig. S5).

**Figure 3.**
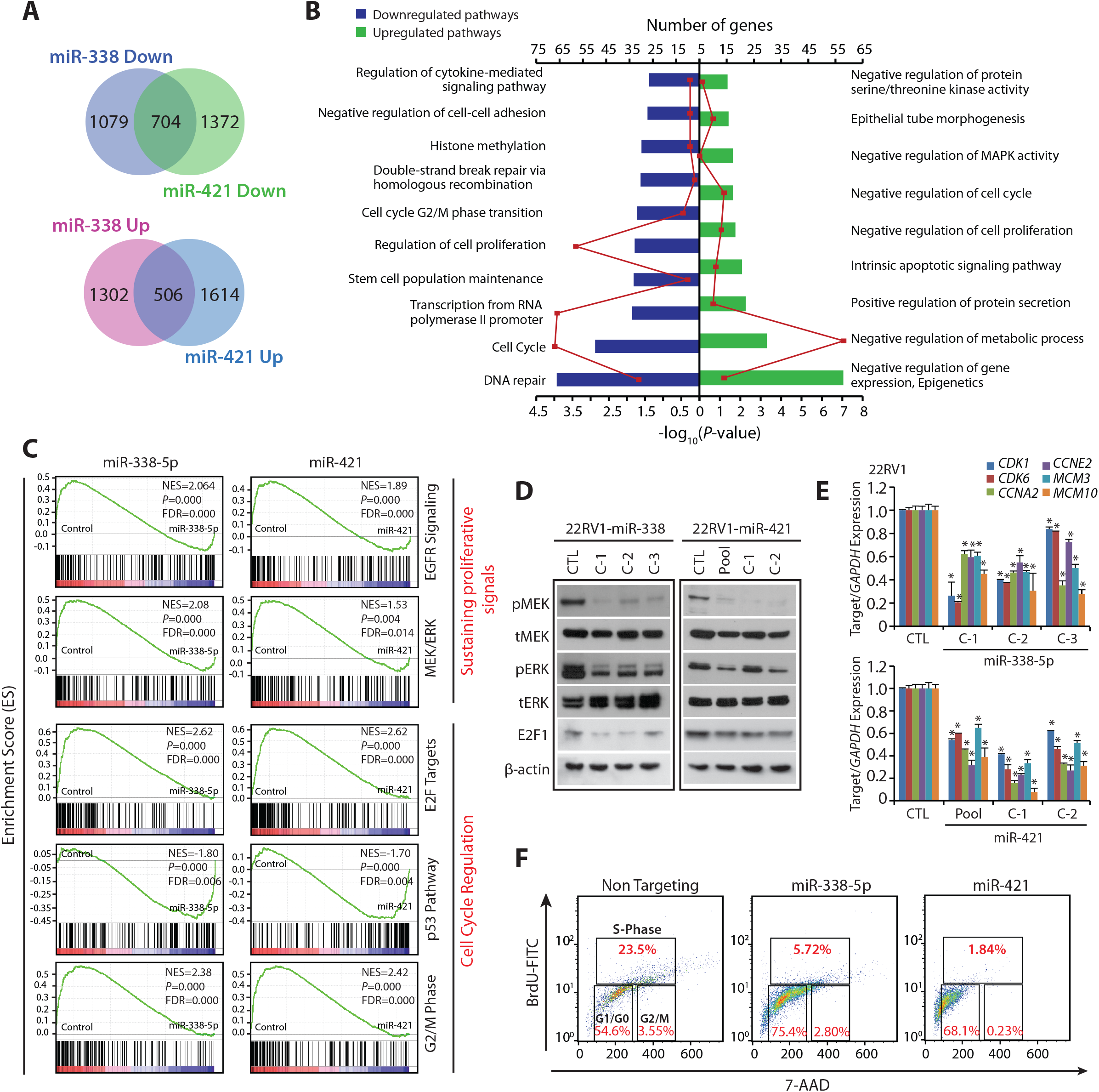
MiR-338-5p and miR-421 overexpression suppress oncogenic pathways and triggers G1/S arrest. **(A)** Gene expression profiling data showing overlap of downregulated (upper panel) and upregulated genes (lower panel) in stable 22RV1-miR-338-5p and 22RV1-miR-421 cells relative to 22RV1-CTL cells (n=3 biologically independent samples). **(B)** Same as in (A), except DAVID analysis showing various downregulated (left) and upregulated (right) pathways. Bars represent –log_10_ (*P*-values) and frequency polygon (line in red) represents the number of genes. **(C)** Gene Set Enrichment Analysis (GSEA) plots showing various deregulated oncogenic gene signatures with the corresponding statistical metrics in the same cells as in (A). **(D)** Western blot analysis for phosphor (p) and total (t) MEK1/2, ERK1/2 and cell cycle regulator E2F1 levels. β-actin was used as a loading control. **(E)** QPCR analysis showing expression of cell cycle regulators for G1 and S phase as indicated. Expression level for each gene was normalized to *GAPDH*. **(F)** BrdU/7-AAD cell cycle analysis for S-phase arrest in 22RV1 cells transfected with miR-338-5p or miR-421 mimics relative to control cells. In the panels (D), (E) and (F) biologically independent samples were used (n=3); data represents mean ± SEM ∗*P*≤ 0.05 and ^∗∗^*P*≤ 0.005 using two-tailed unpaired Student’s *t* test.

Since MAPK signaling pathways involving a series of protein kinase cascades play a critical role in the regulation of cell proliferation, we examined the phosphorylation status of MEK (pMEK) and ERK (pERK), as a read-out of this pathway. In agreement with our *insilico* analysis, a significant decrease in pMEK and pERK was observed in 22RV1-miR-338-5p and 22RV1-miR-421 cells (Fig. 3D). E2F transcription factors are known to interact with phosphorylated retinoblastoma, and positively regulate genes involved in S-phase entry and DNA synthesis (21), thus we next examined the E2F1 level in miRNAs overexpressing cells, surprisingly a notable decrease in E2F1 was observed (Fig. 3D). Further, a significant decrease in the expression of genes involved in G1/S transition such as cyclin E2 (*CCNE2*), cyclin A2 (*CCNA2*) and cyclin-dependent kinase (*CDK1* and *CDK6*), including mini-chromosome maintenance (*MCM3* and *MCM10*), required for the initiation of eukaryotic replication machinery was recorded (Fig. 3E). Thus, these findings corroborate with previous literature that during DNA damage, CDKs being cell-cycle regulators crosstalk with the checkpoint activation network to temporarily halt the cell-cycle progression and promote DNA repair (22). Intriguingly, presence of putative miR-338-5p/miR-421 binding sites on the 3’UTRs of these cell cycle regulators (Supplementary Table S6) further support that these targets could be directly controlled by these miRNAs. Next, to validate that miR-338-5p/-421 overexpression leads to S-phase arrest, 22RV1 cells transfected with miR-338-5p or miR-421 mimics were subjected to cell cycle analysis, a significant increase in the S-phase arrested cells was noted (Supplementary Fig. S6A). To delineate that this increase in the S-phase cells is indeed due to cell-cycle arrest and not because of DNA replication, BrdU-7AAD-based cell cycle analysis was performed, which revealed a significant decrease in the percentage of BrdU incorporated cells in S-phase (Fig. 3F). Taken together, our findings strongly indicate that miR-338-5p/-421 overexpression led to S-phase arrest, thus elucidating the mechanism for reduced cell proliferation and dramatic regression in tumor growth.

### Ectopic expression of miR-338-5p and miR-421 suppresses Epithelial-to-Mesenchymal Transition (EMT) and stemness

Association between EMT and cancer stem cells (CSCs) has been well-established, indicating that a subpopulation of neoplastic cells, which harbor self-renewal capacity and pluripotency, are associated with highly metastatic and drug-resistant cancers (23). Since miR-338-5p/-421 overexpression in 22RV1 cells show regression in tumor burden and metastases (Fig. 2E-G), we evaluated our microarray data for the genes involved in EMT and stemness and noted a marked decrease in their expression (Fig. 4A) including the EMT-inducing transcription factors namely, *SNAI1* (*SNAIL*), *SNAI2* (*SLUG*), and *TWIST1* (Fig. 4B, C). Since, SNAIL and SLUG are known to negatively regulate *CDH1 (E-Cadherin*) (24), an epithelial marker involved in cell-cell adhesion, we next examined E-Cadherin expression. Interestingly, miR-338-5p/-421 overexpressing cells showed a prominent increase in the membrane localization of E-Cadherin, while a significant decrease in the expression of vimentin, a mesenchymal marker was observed (Fig. 4D).

**Figure 4.**
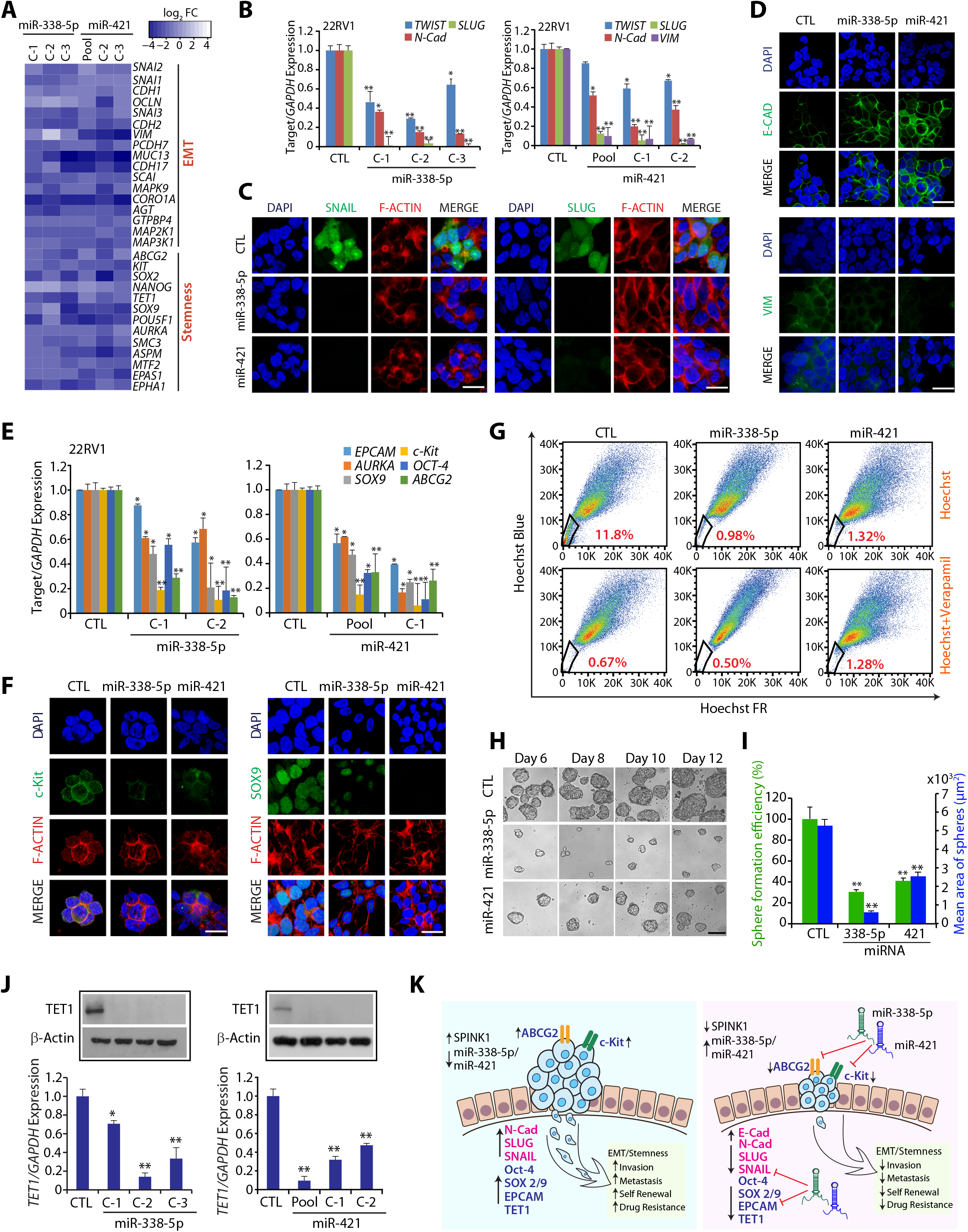
MiR-338-5p and miR-421 overexpression attenuates EMT and Stemness. **(A)** Heatmap depicting change in the expression of EMT and pluripotency markers in 22RV1-miR-338-5p and 22RV1-miR-421 cells. Shades of blue represents log_2_ fold-change in gene expression (n=3 biologically independent samples). **(B)** QPCR analysis depicts expression of EMT markers in 22RV1-miR-338-5p, 22RV1-miR-421 and control cells. Expression for each gene was normalized to *GAPDH*. **(C)** Immunostaining showing SLUG and SNAIL expression in the same cells as in (B). **(D)** Same cells as in (B), except immunostained for E-cadherin and Vimentin. **(E)** Same cells as in (B), except qPCR analysis for stem cell markers. **(F)** Same cells as in (B), except immunostained for c-Kit and SOX-9. **(G)** Hoechst 33342 staining for side population (SP) analysis using same cells as in (B). Percentages of SP were analyzed using the blue and far red filters, gated regions as indicated (red) in each panel. **(H)** Phase contrast microscope images for the prostatospheres using same cells as in (B). Scale bar 100μm. **(I)** Bar plot depicts percent sphere formation efficiency and mean area of the prostatosphere. **(J)** Expression of *TET1* by qPCR and Western blot using same cells as in (B). **(K)** Schematic describing the role of miR-338-5p and miR-421 in regulating EMT, cancer stemness and drug resistance in SPINK1+ cancer. For panels (C), (D) and (F), scale bar represents 20µm. In the panels (B), (E), (I) and (J) biologically independent samples were used (n=3); data represents mean ± SEM ^∗^*P*≤ 0.05 and ^∗∗^*P*≤ 0.005 using two-tailed unpaired Student’s *t* test.

A sub-population (*CD117^+^/ABCG2^+^*) of 22RV1 cells, known as prostate carcinoma-initiating stem-like cells, has been shown to exhibit stemness and multi-drug resistance (25). Therefore, we examined the expression of genes associated with cancer stem cell-like properties in 22RV1-miR-338-5p and 22RV1-miR-421 cells. Strikingly, the expression of well-known pluripotency markers, such as *AURKA, SOX9* and *OCT-4*, and stem-cell surface markers *EPCAM, CD117* (*c-Kit*), and *ABCG2*, an ATP-binding cassette transporter, were markedly downregulated in miRNAs overexpressing cells (Fig. 4E, F). Having confirmed that miR-338-5p/-421 downregulate expression of *ABCG2* and *c-Kit*, we next evaluated the efflux of Hoechst dye *via* ABC-transporters in the absence or presence of verapamil, a competitive inhibitor for ABC transporters (26). As expected, 22RV1-miR-338-5p and 22RV1-miR-421 cells show a significant reduction (~91% and 89% respectively) in the side population (SP) cells involved in Hoechst dye efflux (Fig. 4G). Efflux assay performed in the presence of verapamil show substantial reduction in the SP cells due to inhibition of ABC transporters in both control and miRNAs overexpressing cells (Fig. 4G). Further, to confirm that overexpression of these miRNAs lead to decrease in CSC-like properties, prostatosphere assay, a surrogate model for testing enhanced stem cell-like properties, was performed. As expected, 22RV1-miR-338-5p and 22RV1-miR-421 cells showed a significant decrease in the size and prostatosphere formation efficiency (Fig. 4H, I). Moreover, prostatospheres formed by miRNAs overexpressing cells exhibit a significant reduction in the expression of genes implicated in cancer cells self-renewal and stemness (Supplementary Fig. S6B). Intriguingly, miR-338-5p and miR-421 putative binding sites on the 3’UTR of *EPCAM, c- Kit, SOX9, SOX2* and *ABCG2* were also noticed (Supplementary Table S6), suggesting a possible mechanism involved in the downregulation of these genes.

Epigenetic regulators, such as ten-eleven-translocation (TET) family member, TET1, convert 5’-methylcytosine (5mC) to 5’-hydroxymethylcytosine (5hmC), are well-known to induce pluripotency and maintain self-renewal capacity (27). Thus, we analyzed the expression of TET family members in miRNAs overexpressing cells; strikingly a significant decrease in TET1 was observed (Fig. 4J). Since, *ABCG2* and *c-Kit*, which are implicated in drug-resistance, were downregulated in miRNAs overexpressing cells, the sensitivity of these cells to chemotherapeutic drug was evaluated. Interestingly, 22RV1-miR-338-5p and 22RV1-miR-421 cells show enhanced sensitivity to doxorubicin as compared to control (Supplementary Fig. S6C). Collectively, miR-338-5p/-421 downregulate the expression of genes implicated in multiple oncogenic pathways namely EMT, stemness and drug resistance, signifying that these two tumor suppressor miRNAs could represent a novel approach for integrative cancer therapy (Fig. 4K).

### Transcriptional repression of miR-338 and miR-421 by EZH2 drives SPINK1-positive prostate cancer

Aberrant transcriptional regulation, genomic loss or epigenetic silencing are well-known mechanisms involved in miRNAs deregulation (28,29). Since SPINK1+ PCa patients exhibit reduced expression of miR-338-5p/-421, we sought to decipher the mechanism involved in miRNAs silencing. EZH2, being a member of Polycomb group protein play critical role in epigenetic gene silencing by promoting H3K27me3 marks. Thus, we interrogated Memorial Sloan Kettering Cancer Center (MSKCC) patients’ cohort using cBioPortal (http://cbioportal.org) for any plausible association between *SPINK1* and *EZH2* expression. Interestingly, most of the SPINK1+ specimens comprising Gleason scores 3 and 4 show concordance with EZH2 expression (Fig. 5A). Further, TCGA-PRAD patients harboring higher expression of *EZH2* show increased levels of *SPINK1* and decreased expression of miR-338-5p/-421 as compared to EZH2-low patients (Fig. 5B). To further confirm the association between two oncogenes, we subsequently performed immunohistochemistry (IHC) and RNA in situ hybridization (RNA-ISH) for SPINK1 and *EZH2* expression respectively using tissue microarrays (TMAs) comprising a total of 238 PCa specimens. In accordance with the previous reports (3,4), we also found only 21% PCa patients (50 out of 238 cases) were positive for SPINK1 expression. Interestingly, 88% (44 cases) of these SPINK1+ specimens show positive staining for *EZH2*, of which ~14% of the SPINK1+/EZH2+ patients fall into the high *EZH2* range (score 3 and 4), ~36% in medium *EZH2* (score 2) and ~50% into low *EZH2* expression group (score 1) indicating a significant association between SPINK1 and *EZH2* expression (Fig. 5C; χ^2^=13.66; *p*=0.008). Although, 75% (141 cases) of the SPINK1-negative (SPINK1–) patients were also found positive for *EZH2*, while majority of these cases (~71%) exhibit low *EZH2* levels (score 1). Thus, in corroboration with the previous reports (1,10), our data indicate a more pronounced role of epigenetic alterations involved in *ETS*-fusion negative prostate cancer. Although, six SPINK1+ cases failed to show any expression of EZH2, pointing that an alternative mechanism may be involved in SPINK1 regulation or possibly miRNA-338/-421 genomic deletion could be a cause.

**Figure 5.**
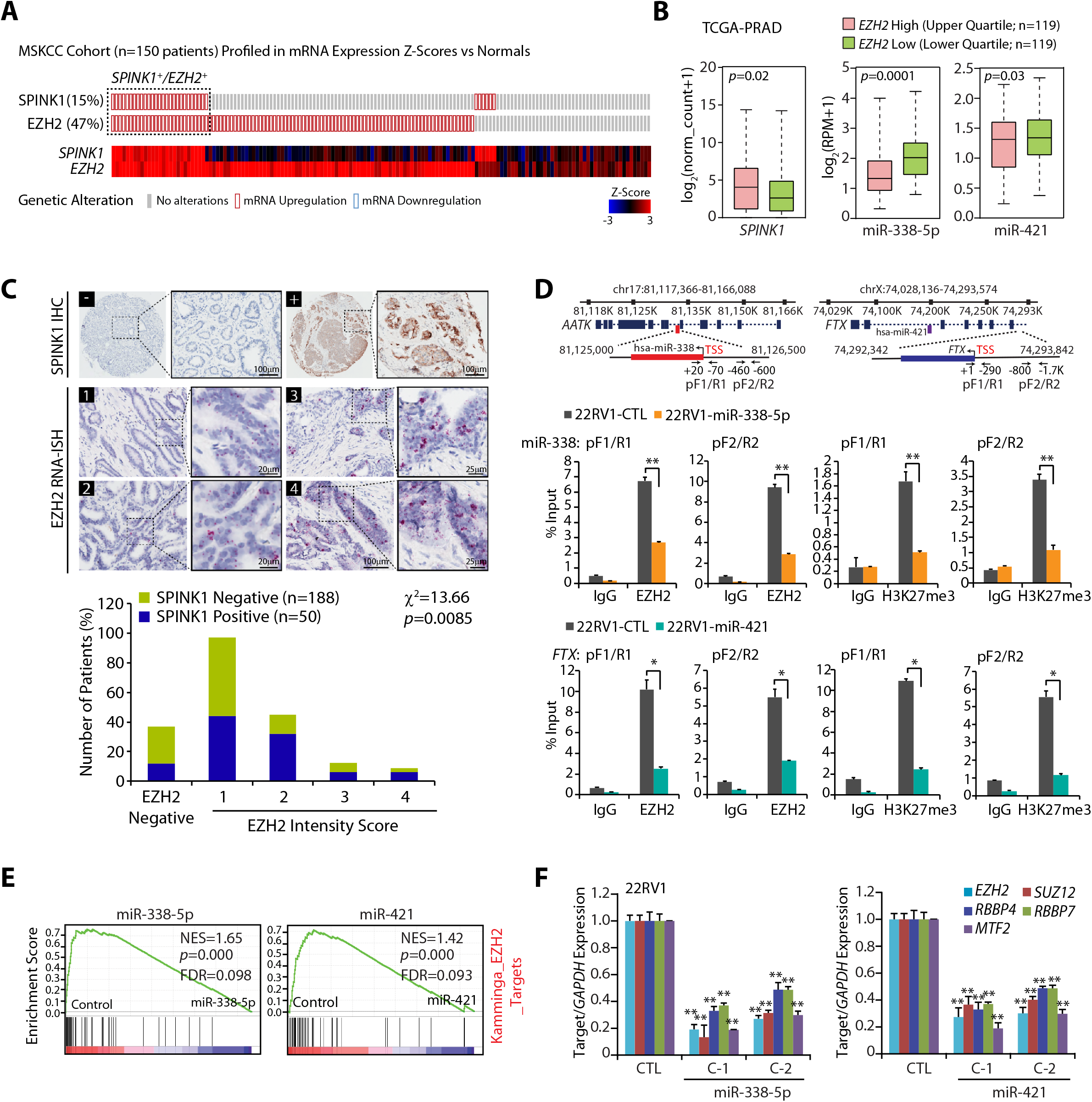
Epigenetic silencing of miR-338-5p and miR-421 via EZH2 in SPINK1 positive prostate cancer. **(A)** OncoPrint depicting mRNA upregulation of *EZH2* and *SPINK1* in MSKCC cohort using cBioportal. In the lower panel shades of blue and red represents Z-score normalized expression for *EZH2* and *SPINK1*. **(B)** Box plot depicting *SPINK1*, miR-338-5p and miR-421 expression in *EZH2* high (n=119) and *EZH2* low (n=119) in PCa patients from TCGA-PRAD cohort **(C)** Representative micrographs depicting PCa tissue microarray (TMA) cores (n=238) stained for SPINK1 by immunohistochemistry (IHC) and *EZH2* by RNA in-situ hybridization (RNA-ISH). Top panel represents SPINK1 IHC in SPINK1 negative (−) and SPINK1 positive (+) patients. RNA-ISH intensity score for *EZH2* expression was assigned on a scale of 0 to 4 according to visual criteria for the presence of transcript at 40X magnification. Bar plot show EZH2 expression in the SPINK1-negative and SPINK1+ patient specimens. *P*-value for Chi-square test is indicated. **(D)** Genomic location for EZH2 binding sites on the miR-338 and *FTX* promoters and location of ChIP primers (top panel). ChIP-qPCR data showing EZH2 occupancy and H3K27me3 marks on the miR-338, *FTX* promoters, and *MYT1* used as positive control in stable 22RV1-miR-338-5p, 22RV1-miR-421 and 22RV1-CTL cells. **(E)** GSEA plots showing the enrichment of EZH2 interacting partners (Kamminga) in 22RV1-miR-338-5p and 22RV1-miR-421 cells. **(F)** QPCR data showing expression of *EZH2* and its interacting partners in the same cells as indicated. Biologically independent samples (n=3) were used in panels (D) and (F); data represent mean ± SEM. ^∗^*P*≤ 0.05 and ^∗∗^*P*≤ 0.005 using two-tailed unpaired Student’s *t* test.

To investigate whether epigenetic silencing of these miRNAs is mediated by EZH2, we screened the promoters of miR-338, miR-421 and *FTX* (miR-421 host gene) for putative transcription factor binding sites and identified MYC and MAX (Myc-Associated Factor X) elements within ~2 kb upstream of Transcription Start Site (TSS). MYC is known to form a repressive complex with EZH2 and HDACs, and downregulate multiple tumor suppressive miRNAs, which in turn target PRC2-interacting partners (30). In addition, EZH2-silenced DU145 cells miRNA expression data (GSE26996) indicates an increase in the expression of numerous EZH2-regulated miRNAs including miR-338 and miR-421 (Supplementary Fig. S7A). We therefore examined the promoters of miR-338 and *FTX* for the recruitment of EZH2 in stable 22RV1-miR-338-5p, 22RV1-miR-421 and control cells. Interestingly, a significant enrichment of EZH2 over input was observed on the promoters of miR-338 and *FTX* in control cells, while substantial decrease in miR-338-5p/-421 overexpressing cells was noticed (Fig. 5D), suggesting presence of negative feedback regulatory network between miR-338-5p/-421 and EZH2. Subsequently, we also checked for EZH2 recruitment on miR-421 promoter and no enrichment was observed (Supplementary Fig. S7B), indicating that the host gene *FTX* promoter regulates the expression of this intronic miRNA. Further, to confirm EZH2-mediated methyltransferase activity, we sought to identify H3K27me3 marks on these promoters. A remarkable enrichment of H3K27me3 marks on the miR-338, miR-421 and *FTX* promoters were noted relative to IgG control (Fig. 5D and Supplementary Fig. S7B), confirming the role of EZH2 mediated epigenetic silencing of miRNA-338-5p/-421.

Comprehensive GSEA analysis revealed that miRNA-338/-421 overexpressing cells show an enrichment for EZH2 interacting partners, including PRC2 members (Kamminga *et al*., 2006) and EZH2 regulated genes (Lu *et al*., 2010; Nuytten *et al*., 2008) (Fig. 5E and Supplementary Fig. S7C), indicating that these two miRNAs in turn regulate EZH2 partners and their target genes. Thus, we next examined the putative binding of miRNA-338-5p/-421 on the 3’UTR of the PRC2 members, interestingly both miRNAs show negative mirSVR binding score (Supplementary Table S6). Moreover, a significant decrease in the transcript levels of *EZH2*, and its interacting partners *SUZ12, RBBP4, RBBP7* and *MTF2* were observed in miRNA-338-5p/-421 overexpressing cells (Supplementary Fig. S7D and Fig. 5F). Collectively, our data indicates that overexpression of miR-338-5p/-421 downregulates EZH2 expression and its interacting members, leading to impaired histone methyltransferase activity of PRC2, thereby establishing a double-negative feedback loop.

Since inhibitors for chromatin modifiers are known to erase epigenetic marks, we tested 3-Deazaneplanocin A (DZNep), an inhibitor of the histone methyltransferase; 2’-deoxy-5-azacytidine (5-Aza), a DNA methyltransferase (DNMT) inhibitor and Trichostatin A (TSA), a HDAC inhibitor in 22RV1 cells and examined the expression of miR-338-5p/-421. Treatment with TSA, DZNep, 5-Aza alone or a combination of DZNep and TSA in 22RV1 cells showed a modest increase in miR-338-5p/-421 expression, while 5-Aza and TSA together resulted in ~9-fold increase (Fig. 6A). Intriguingly, 5-Aza and TSA in combination results in significant increase in miRNAs expression accompanied with a notable decrease (~60-80%) in *SPINK1* levels (Fig. 6B). Since, 3’-arm of miR-338 (miR-338-3p) is known to negatively regulate *Apoptosis Associated Tyrosine Kinase (AATK*) expression (31), likewise a significant reduction in the *AATK* expression was noticed in our study (Fig. 6B). Furthermore, a deletion construct of *FTX* show decreased expression of miR-374/-421 cluster (32). In line with this, a significant increase in the *FTX* and miR-421 expression was reported upon 5-Aza and TSA combinatorial treatment, signifying the importance of host gene *FTX* in the regulation of miR-421 (Fig. 6B).

**Figure 6.**
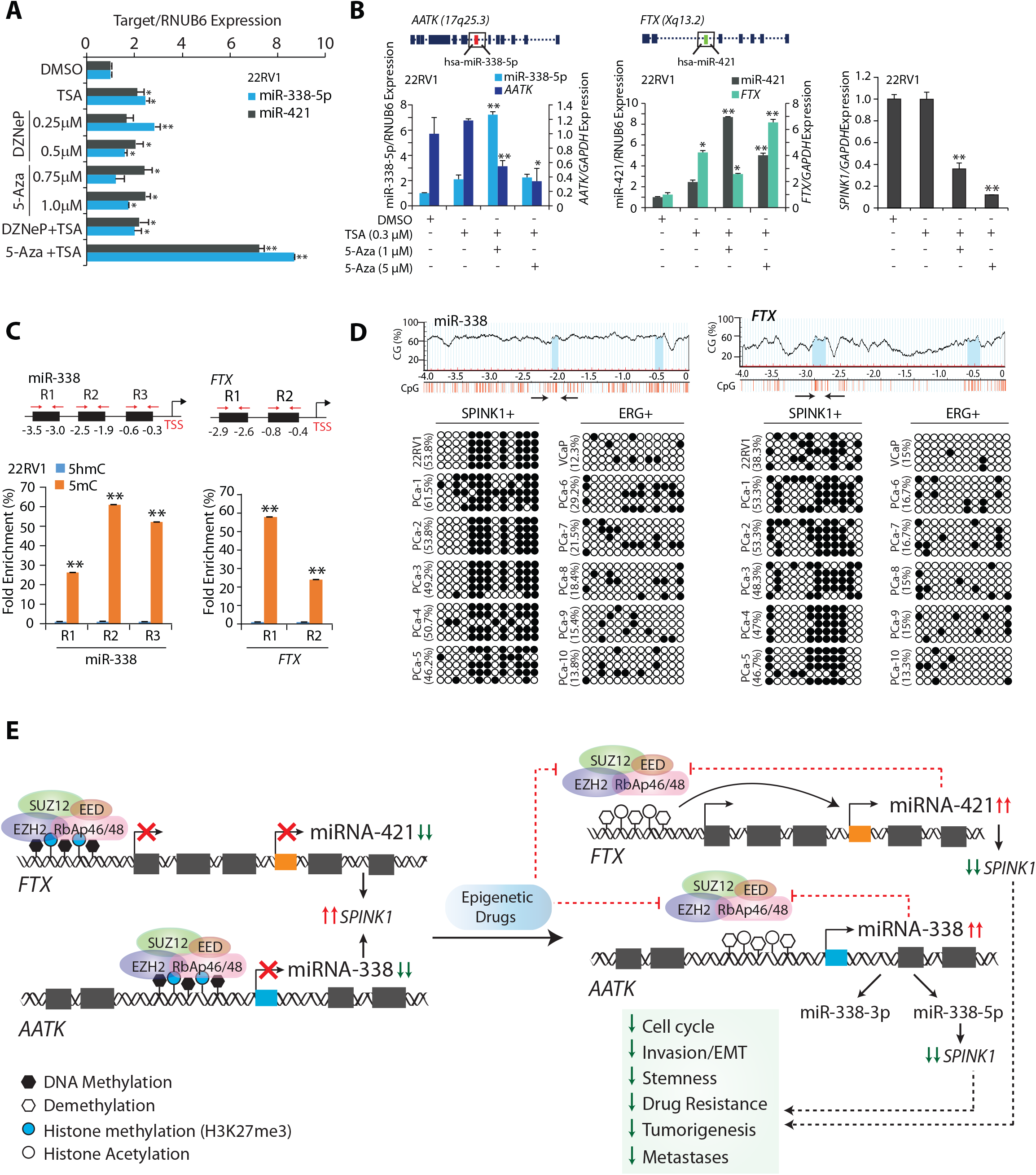
Epigenetic drugs ablate EZH2-mediated silencing of the miR-338-5p and miR-421. **(A)** TaqMan assay for miR-338-5p and miR-421 expression in 22RV1 cells treated with different combination of epigenetic drugs. **(B)** QPCR showing relative expression of miR-338-5p, miR-421, *AATK*, *FTX* and *SPINK1* in 22RV1 cells treated with 5-Aza or TSA as indicated. **(C)** MeDIP-qPCR showing fold enrichment of 5-mC over 5-hmC in 22RV1 cells as indicated. **(D)** Bisulfite–sequencing showing CpG methylation marks on the region upstream of miR-338-5p (left) and *FTX* (right) in 22RV1, VCaP cells and patients’ tumor specimens (PCa-1 to 5 are SPINK1 positive and PCa-6 to 10 are ERG fusion positive). PCR amplified regions are denoted by arrows. Data represents DNA sequence obtained from five independent clones. Hollow circles represent nonmethylated CpG dinucleotides, whereas black solid circles show methylated-CpG sites. **(E)** Illustration depicting the molecular mechanism involved in EZH2-mediated epigenetic silencing of miR-338-5p and miR-421 in *SPINK1*-positive prostate cancer. In panels (A), (B) and (C) biologically independent samples (n=3) were used; data represent mean ± SEM. ∗*P*≤ 0.05 and ^∗∗^*P*≤ 0.005 using two-tailed unpaired Student’s *t* test.

In addition, EZH2 is also known to interact with DNMTs, thus enabling chromatin remodeling and DNA methylation (33). Hence, we next examined the presence of methylated CpG marks on the promoters of miR-338-5p and *FTX*. Interestingly, methylated DNA immunoprecipitation (MeDIP) revealed locus-specific enrichment in the 5mC levels over 5hmC on these regulatory regions (Fig. 6C). To ascertain the presence of DNA methylation marks we performed bisulfite sequencing using PCa cell lines, a relative increase in the methylated CpG sites on miR-338 and *FTX* promoters was observed in 22RV1 cells (SPINK1-positive) as compared to VCaP (ERG-positive) cells (Fig. 6D). No significant difference in the methylated CpG sites on the *AATK* and miR-421 promoters was observed (Supplementary Fig. S7E, F). To understand clinical relevance, bisulfite sequencing was carried out on SPINK1-positive (n=5) and *ERG* fusion positive (n=5) PCa patients’ specimens. Interestingly, all SPINK1-postive specimens exhibit increased methylation marks on the promoters of miR-338 and *FTX* as compared to *ERG* positive (Fig. 6D). Taken together, our results strongly indicate that epigenetic machinery comprising of EZH2 and its interacting partners play a critical role in the epigenetic silencing of miRNA-338-5p and miR-421 in *SPINK1+* subtype, which in turn reaffirms its silencing by a positive feedback loop.

## Discussion

In this study, we unraveled the underlying molecular mechanism involved in the overexpression of SPINK1 exclusively in *ETS*-fusion negative PCa. Our study provides a molecular basis for SPINK1 overexpression, brought about by the epigenetic repression of key post-transcriptional negative regulators of *SPINK1* namely, miR-338-5p and miR-421. We demonstrated the tumor suppressive roles of miR-338-5p/-421, which exhibits functional anti-cancer pleiotropy in *SPINK1+* subtype, by attenuating oncogenic properties, tumor growth and metastases in murine model. Conversely, miR-421 has also been reported to be a potential oncogenic miRNA in multiple cancers (34,35). However, in corroboration with our findings, a recent report suggested tumor suppressive role of miR-421 in prostate cancer (36). We also established that miR-338-5p/-421 overexpressing cells display perturbed cell-cycle machinery triggered by dysregulated cyclins and CDKs, subsequently leading to S-phase arrest. It has been shown that miRNAs targeting multiple cyclins/CDKs are more effective than the FDA-approved CDK4/6 inhibitor in triple-negative breast cancer (37), thus supporting our findings that replenishing these miRNAs may prove advantageous in SPINK1+ cancers.

Emerging evidences suggest a complex interaction between EMT and CSCs during cancer progression, and in developing resistance towards anti-cancer drugs. Previous studies have implicated the role of several miRNAs, such as miR-200 family, miR-205 and miR-34a (38,39) in regulating the expression of genes involved in metastases, stemness and drug resistance. Furthermore, miR-338 exhibits tumor suppressive role, and inhibits EMT by targeting *ZEB2* (40) and *PREX2a* (41) in gastric cancer. Here, we identified miR-338-5p/-421 as critical regulators of EMT-inducing transcription factors and -associated markers, which in turn led to decreased stem-cells like features. Moreover, CSCs are known to express ABC transporters, which efflux the chemotherapeutic drugs during resistance (26). Remarkably, miR-338-5p/-421 overexpression shows decreased expression of *ABCG2* and *c-KIT*, consequently a significant drop in the drug-resistant side population, indicating that these two miRNAs are highly effective in conferring drug-sensitivity and reducing the therapy-resistant CSCs. Collectively, our findings provide a solid foundation for qualifying these miRNAs as an adjuvant therapy for the *SPINK1+* as well as other drug resistant malignancies.

These findings were further strengthened by a recent TCGA study (1), wherein a subset of PCa patients’ harboring *SPOP*-mutation/*CHD1*-deletion exhibits elevated DNA methylation levels accompanied with frequent events of *SPINK1* overexpression. Recently, a new subtype of *ETS*-fusion-negative tumors has been defined by frequent mutations in the epigenetic regulators and chromatin remodelers (42). Yet another study, using genome-wide methylated DNA-immunoprecipitation sequencing revealed higher number of methylation events in *TMPRSS2-ERG* fusion-negative as compared to normal and *TMPRSS2–ERG* fusion-positive PCa specimens (10), thus collectively, these independent findings reaffirm the critical role of epigenetic pathways engaged in the pathogenesis of *SPINK1+* subtype.

Interestingly, increased methylated regions in the *ETS*-fusion negative patients have been attributed to hypermethylation of miR-26a, a post-transcriptional regulator of EZH2 (10). Thus, given the central role played by EZH2 and the epigenetic mechanism involved in *ETS*-fusion negative cases, our findings rationalize the role of EZH2-mediated epigenetic regulation of miR-338-5p and miR-421 in *SPINK1+*/*ETS*-negative subtype. Hence, we propose a molecular model involving *SPINK1*, *EZH2* and miR-338-5p/-421, wherein EZH2 acts as an epigenetic switch and by its histone methyltransferase activity establishes H3K27me3 repressive marks on the regulatory regions of miR-338 and *FTX*, a miR-421 host gene (Fig. 6E).

In consonance with this, overexpression of miR-338-5p/-421 also results in decreased Tet1 expression. Converging lines of evidences suggest dual role of Tet1 in promoting transcription of pluripotency factors as well as recruitment of PRC2 on the CpG rich promoters (43). Taken together, miR-338-5p/-421 mediated decrease in Tet1 expression might possibily contribute in reduced stemness and drug-resistance. We also conjecture that decrease in Tet1 expression may result in reduced PRC2 occupancy on the miRNA promoters, diminish epigenetic silencing marks, and consequently downregulate their targets including *SPINK1*.

Currently, there is no effective therapeutic intervention for *SPINK1*+/*ETS*-negative PCa as well as for other SPINK1+ malignancies, although use of monoclonal EGFR antibody has been suggested (8). Nevertheless, outcome of the phase I/II clinical trials using cetuximab (44) and small molecules inhibitors for EGFR has been largely unsuccessful (45,46). For instance, in a phase Ib/IIa clinical trial using cetuximab and doxorubicin combination therapy, only a fraction of CRPC patients (~8%) showed >50% PSA decline (44), revealing its limited efficacy. Owing to the pleiotropic anti-cancer effects exhibited by miRNA-338-5p/-421, we propose miRNA-replacement therapy as one of the potential therapeutic approaches for SPINK1+ cancers; nonetheless *in-vivo* delivery methods and stability are some of the major challenges for successful translation into the clinic (47). While not restricted to miRNA-replacement therapy, the present study also suggests alternative therapeutic avenues for *SPINK1+* malignancies, for instance adjuvant therapy using inhibitors against DNMTs, HDACs or EZH2, several of which are already in clinical trials (48). Conclusively, we moved the field forward by addressing an important question that how SPINK1 is aberrantly overexpressed in *ETS*-fusion negative prostate cancer, and the stratification of patients based on *SPINK1*-positive and miRNA-338-5p/-421-low criteria could further improve therapeutic modalities and overall management strategies.

## Data availability

The gene expression microarray data from this study has been submitted to the NCBI Gene Expression Omnibus (GEO, http://www.ncbi.nlm.nih.gov/geo/) under the accession number GSE108558.

## Acknowledgements

B.A. is an Intermediate Fellow of the Wellcome Trust/DBT India Alliance. This work is supported by the Wellcome Trust/DBT India Alliance Fellowship [grant number: IA/I(S)/12/2/500635] awarded to B.A. We thank Yuping Zhang, Brendan Veeneman, Mahendra Palecha, Ayush Praveen for their technical support and Anjali Bajpai for critically reading the manuscript. We also thank Jonaki Sen for extending the use of fertilized eggs facility. The IIT Kanpur has filed a patent (IN 201611016564) on the therapeutic applicability of miR-338-5p and miR-421 described in this study in which B.A., V.B. and A.Y. are named as inventors.

## Authors’ contributions

V.B., A.Y., and B.A. designed and directed the experimental studies. V.B. and A.Y. performed *in-vitro* cell line-based studies. V.B., A.Y. and S.N. performed the bisulfite sequencing experiments and analysis. V.B., A.Y., R.T. and S.G. performed the gene expression studies, bioinformatics analysis and ChIP assays. A.Y. performed the immunofluorescence and RIP experiments. V.B., A.Y., and B.A. performed statistical analysis and interpreted the data. V.B., A.Y., and B.A. executed the *in-vivo* experiments. A.G. provided PCa patient specimens. S.C., N.G., and N.P. performed immnuohistochemistry and RNA *in situ* staining on the PCa tissue microarrays. V.B., A.Y., and B.A. wrote the manuscript. B.A. directed the overall project.

